# Genome-resolved metagenomic analyses reveal the presence of a bacterial endosymbiont in an avian nasal mite (Rhinonyssidae; Mesostigmata)

**DOI:** 10.1101/2021.07.12.452008

**Authors:** Carolina Osuna-Mascaró, Jorge Doña, Kevin P. Johnson, Manuel de Rojas

**Affiliations:** Department of Biology, University of Nevada, Reno; Illinois Natural History Survey, Prairie Research Institute, University of Illinois at Urbana-Champaign, Champaign, USA; Departamento de Biología Animal, Universidad de Granada, Granada, Spain; Department of Microbiology and Parasitology, Faculty of Pharmacy, Universidad de Sevilla, Sevilla, Spain

**Keywords:** Rhinonyssidae, endosymbiont, metagenomic, Brucella

## Abstract

Rhinonyssidae (Mesostigmata) is a family of nasal mites only found in birds. All species are hematophagous endoparasites, which may damage the nasal cavities of birds, and also could be potential reservoirs or vectors of other infections. However, the role of members of Rhinonyssidae as disease vectors in wild bird populations remains uninvestigated, with studies of the microbiomes of Rhinonyssidae being almost non-existent. In the nasal mite (*Tinaminyssus melloi*) from rock doves (*Columba livia*), a previous study found evidence of a highly abundant putatively endosymbiotic bacteria from Class Alphaproteobacteria. Here, we expanded the sample size of this species, incorporated contamination controls, and increased sequencing depth in shotgun sequencing and genome-resolved metagenomic analyses. Our goal was to increase the information regarding this mite species with its putative endosymbiont. Our results support the endosymbiotic nature of this bacterial taxon, which is the first described for bird’s nasal mites to date, and improve the overall understanding of the microbiota inhabiting these mites.

## 1. Introduction

Mites are one of the most diverse groups of eukaryotes on earth [1]. Mites are ubiquitous, occupying aquatic, terrestrial, and arboreal habitats [2–8]. On some occasions, mites have an intimate association (symbiosis) with a different organism, which, in the most extreme scenario (permanent symbionts), represents the habitat in which they undergo their entire life-cycle [9]. For example, a single species of insect, mammal, or bird can often host several mite species and have different types of interactions with them [9, 10–12]. The nature of these interactions can range from mutualism (e.g., mites inhabiting and cleaning birds’ feathers; [13]) to parasitism (e.g., mites inhabiting the nasal passages and lungs of seals causing illness to them; [14, 15]).

In some cases, mites can affect the host’s health and be responsible for transmitting zoonotic diseases [9]. For example, the itch mite *Sarcoptes scabiei* has caused the population decline of different mammal species by transmitting scabies [16–19]. In another case, *Varroa destructor*, a parasitic mite of the Asian honeybee (*Apis cerana*) has caused the decline of the European honeybee by transmitting a virus [20]. In some cases, disease transmission may be mediated by endosymbiotic pathogens that inhabit the mites. One example of this mechanism is *Leptotrombidium scutellare*, a mite that parasitizes mice and carries the endosymbiont bacteria *Orientia tsutsugamushi*, responsible for the scrub typhus disease [21]. Endosymbionts as causal agents of diseases have also been widely reported in ticks [22–24]. However, examples of endosymbionts in mites are still rare.

Rhinonyssidae (Mesostigmata) is a family of nasal mites with more than 500 species described worldwide [25–28]. They have been described from birds [29]. Almost all species of birds are inhabited by nasal mites, which usually live in the nasal cavity on vascularized epithelial tissue [9]. All rhinonyssid species are hematophagous endoparasites [30]. Specifically, rhinonyssid mites damage the nasal cavities of birds, which, in rare cases, may lead to the death of the hosts (Rhinonyssidosis avium disease) [28]. Moreover, it has been suggested that these mites could be potential reservoirs or vectors of other infections (such as West Nile fever, Q fever, avian influenza, and Lyme diseases), as demonstrated in mites of the family Dermanyssidae [31]. However, the role of species of Rhinonyssidae as disease vectors in wild bird populations is yet to be understood. In addition, studies on rhinonyssid microbiomes more generally are almost non-existent.

In a preliminary previous study, the microbiome of two different species of rhinonyssid mites (*Tinaminyssus melloi* and *Ptilonyssus chlori)* from two different avian host species (*Columba livia* and *Chloris chloris*, respectively) was characterized [32]. The results of that study suggested that the nasal mite *Tinaminyssus melloi* harbored a potential endosymbiont Alphaproteobacteria (Family: Bartonellaceae) in a high abundance. However, the low sequencing coverage, small sample size, and lack of control samples did not allow definitive conclusions to be made. In particular, only a partial (~26%) assembly of the bacterial genome was achieved, and it could not be ruled out completely if this bacterial taxon came from the bird host instead of the mite.

Here, we focused on expanding knowledge of the microbiome of this mite species (*Tinaminyssus melloi)*, emphasizing increasing the information of its association with this putative endosymbiont. In the current study, we expanded the sample size, used contamination controls, and increased sequencing depth. In particular, we collected two mite pools (five mite individuals per pool) plus a control bird saliva sample from two different individual Rock doves (*Columba livia)*. Then, we conducted shotgun and genome-resolved metagenomic analyses to characterize the mite’s microbiome along with evaluating the genome properties of this bacterial taxon, which may be informative regarding its endosymbiotic status.

## 2. Materials and Methods

A total of ten nasal mites were collected from two different freshly dead *Columba livia* individuals (i.e., 5 mites per individual host). The nasal cavities of the birds were dissected under a stereomicroscope, and the mites were taxonomically identified. A saliva sample was also collected from each individual. Mite and saliva samples were preserved at −20°C in tubes with 100% ethanol.

Before DNA isolation, samples were washed three times with ethanol to remove possible external contaminants following [13] and [33]. Total genomic DNA was isolated from all samples using the Quick-DNA MicroPrep kit (Zymo), specifically designed to isolate DNA from small samples. A sample that did not contain tissue was included and treated as a regular sample to check for cross-contamination during the DNA isolation procedure. Total DNA was quantified using the Qubit High Sensitivity dsDNA Assay (Thermo Fisher Scientific).

Libraries were prepared using the Nextera DNA Flex Library Prep kit (Illumina), strictly following the manufacturer’s instructions. Briefly, DNA was enzymatically cut, and adapters were added in a single step. The ligated DNA was amplified, and oligonucleotide indices were added to both ends of the fragments for post-sequencing demultiplexing. The constructed libraries were quantified with the Qubit dsDNA HT Assay kit (Thermo Scientific), and quality checked on an Agilent 2100 Bioanalyzer (Agilent Technologies). According to the Qubit results, the libraries were pooled in equimolar amounts, and this pool was sequenced on a NovaSeq PE150 single lane fraction (Illumina), aiming for a total output of 30 gigabases. The DNA isolation, amplification, library preparation, and whole-genome sequencing were carried out in AllGenetics & Biology SL (www.allgenetics.eu).

For the genome-resolved metagenomic analyses, we trimmed the raw reads using fastp [34]. We used BBNorm [35] to reduce the coverage of the concatenated FASTQ file to a maximum of 60X and discarding reads with a coverage under 5X. Using this normalized coverage, we ran the metaWRAP v1.1.5 pipeline [36]. First, we used the metaWRAP Read_qc module with default parameters to quality trim the reads. We then assembled the reads using the metaWRAP Assembly module (–use megahit option) [37]. We binned the reads with the metaWRAP Binning module (–maxbin2 –concoct –metabat2 options), and after that we consolidated the resulting bins into a final bin set with Bin_refinement module (-c 50 -× 10 options). We quantified the bins with the Quant_bin module and then reassembled the consolidate bins set using the Reassemble_bins module. Finally, we classified bins using the Classify_bins module. In addition, we uploaded our final metagenome-assembled genomes (MAGs) to MiGA [38] for a complementary analysis to determine the most likely taxonomic classification and novelty rank of the bin. We used the NCBI Genome database (NCBI Prok; Apr-23-2021 version) for this analysis and the TypeMat database ([38]; r2021-04 version). Additionally, we conducted an investigation of the metabolic capabilities of the assembled bacteria by investigating the completeness of metabolic pathways using GhostKOALA [39] and KEGG-Decoder [40].

For the shotgun metagenomic analyses, we used the metagenomic classifier Kaiju [41] to characterize the taxonomic content of the metagenomes with the following parameters: Reference database: nr +euk; Database date: 2017-05-16; SEG low complexity filter: yes; Run mode: greedy; Minimum match length: 11; Minimum match score: 75; Allowed mismatches: 5. We then converted Kaiju’s output files into a summary table at the genus and species level and filtered out taxa with low abundances (<0.3% of the total reads). We also removed poorly identified taxa because they would artificially increase the similarity between our samples. Specifically, the following taxa were excluded: “NA,” “Viruses,” “archaeon,” “uncultured bacterium,” “uncultured Gammaproteobacteria bacterium”.

To visualize similarities of microbiome composition among mite individuals from different individual hosts and saliva samples, we constructed non-metric multidimensional scaling (NMDS) ordinations based on Bray–Curtis and Jaccard (binary = T) dissimilarities using phyloseq v1.26-1 R package [42]. Prior these analyses, matrices were rarefied using the rarefy_even_depth function of phyloseq (without replacement as in the hypergeometric model) to rarefy samples to the smallest number of classified sequences per individual observed.

Endosymbionts genomes have been typically described as small (i.e., eroded with respect to non-endosymbiotic species) and have an AT base compositional bias [43]. Accordingly, we explored the relative position of the putative endosymbiont MAG in a “Genome size ~ GC content” correlation plot. Specifically, we compared our results to those from Doña et al. 2021 [44], who used this approach to identify potential endosymbionts in lice. Finally, we aligned the MAG to the partial one previously assembled in Osuna-Mascaró et al., 2020 [32], using Mauve (MCM algorithm and default parameters) [45], and we estimated the pairwise genetic distances between both MAGs.

## 3. Results

Rhinonyssid mites from two hosts as well that two saliva samples were sampled and sequenced (see **Table S1** for details). From the genome-resolved metagenomic pipeline, we retrieved a single bacterial metagenome-assembled genome (MAG). The MAG was present in all but the saliva samples (mean MAG copies per million reads: mite samples = 331.5; saliva samples = 0). According to CheckM, the MAG was 98.1 % complete, with only 0.9 % contamination, N50 = 101,044 bp. The MAG has a 99.5 % similarity with the MAG assembled in [32] Osuna-Mascaró et al., 2020 (**Figure 1**). Also, the MAG has characteristics typical of endosymbionts with an AT bias (GC content = 0.31) and a small genome size (1,269,226 bp). In particular, compared to the MAGs from Doña et al. 2021 [44], the MAG has lower GC content, and a smaller genome size than any of the MAGs from that study. The metaWRAP Classify_bins module classified the MAG as belonging to the family Bartonellaceae. A further taxonomic classification analysis in MiGA revealed similar results. When using the TypeMat database, it was classified as possibly belonging to the class Alphaproteobacteria (P = 0.0012) and close to the family Brucellaceae (P = 0.41). Specifically, the closest related species found were *Brucella abortus* (544 GCA 000369945T; 53.03 % AAI, i.e., maximum average amino acid identity) and *Brucella microti* (CCM 4915 NC 013119; 99.73 % AAI). MiGA results indicate that the MAG most likely belongs to a species not represented in the TypeMat database (P = 0.00034) and probably belongs to a genus not represented in the database (P = 0.171). When using the NCBI Prok database, it was also classified as an Alphaproteobacteria (P = 0.0015), and the closest relatives found in this database were *Candidatus Tokpelaia hoelldoblerii* (CP017315; 51.01% AAI) and *Brucella pinnipedialis* (B2 94 NC 015857; 50.71% AAI). Using this database, MiGA analyses also indicate that this species most likely belongs to a species not represented in the database (P = 0.00017) and probably to a genus not represented in the database (P = 0.12).

**Figure 1.**
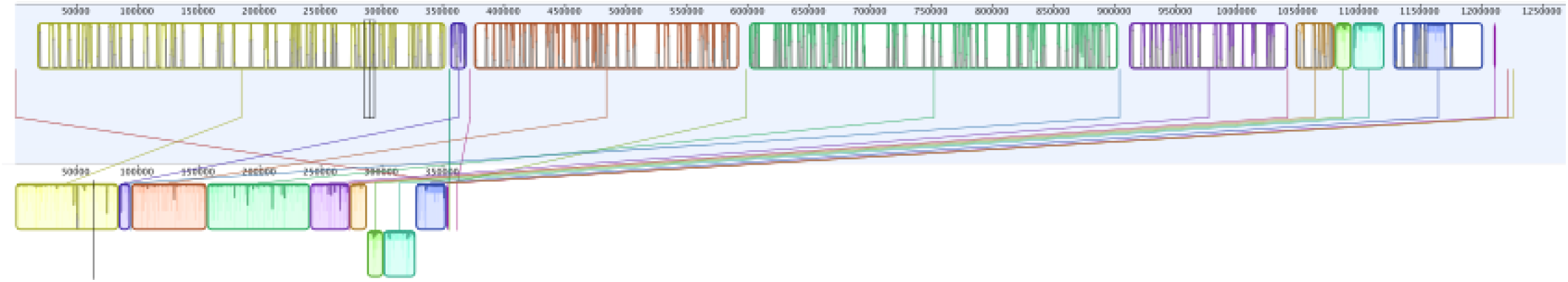
Mauve alignment of the MAG found in this study (on the top; 98.1 % completeness) to the MAG assembled in Osuna-Mascaró et al., 2020 [32] (on the bottom, 26 % completeness).

Lastly, the MAG was classified by NCBI Prok also as a possible Alphaproteobacteria (P = 0.0015), and the closest relative found in this database were *Candidatus Tokpelaia hoelldoblerii* (CP017315; 51.01% AAI) and *Brucella pinnipedialis* (B2 94 NC 015857; 50.71% AAI). We found that this MAG has complete pathways for vitamin B (riboflavin) and vitamin B12 synthesis, among others (**Table S2** and **Figure S1**). In addition, the MAG has complete pathways for synthesis of essential amino acids (i.e., lysine) and several non-essential amino acids (i.e., aspartate, glutamate, serine). We also found many fully or partially missing pathways that may be redundant or potentially shared (or synthesized along) with the mite.

Kaiju analyses recovered a higher diversity of microorganisms. When collapsing the Kaiju matrices at the genus level, the bacterial taxa with higher relative abundances were *Staphylococcus*, *Streptomyces*, *Nocardia*, *Clostridioides*, *Chlamydia*, and *Bartonella* (**Figure 2**). When collapsing at the species level, the bacterial taxa with higher relative abundance were *Staphylococcus capitis*, *Streptomyces shenzhenensis*, *Nocardia nova*, and *Clostridum difficile* (**Figure 3**). NMDS ordinations showed a similar clustering among mite samples and saliva samples when collapsing at both the species and the genus level, and based on both Bray–Curtis and Jaccard distances (**Figure S2**).

**Figure 2.**
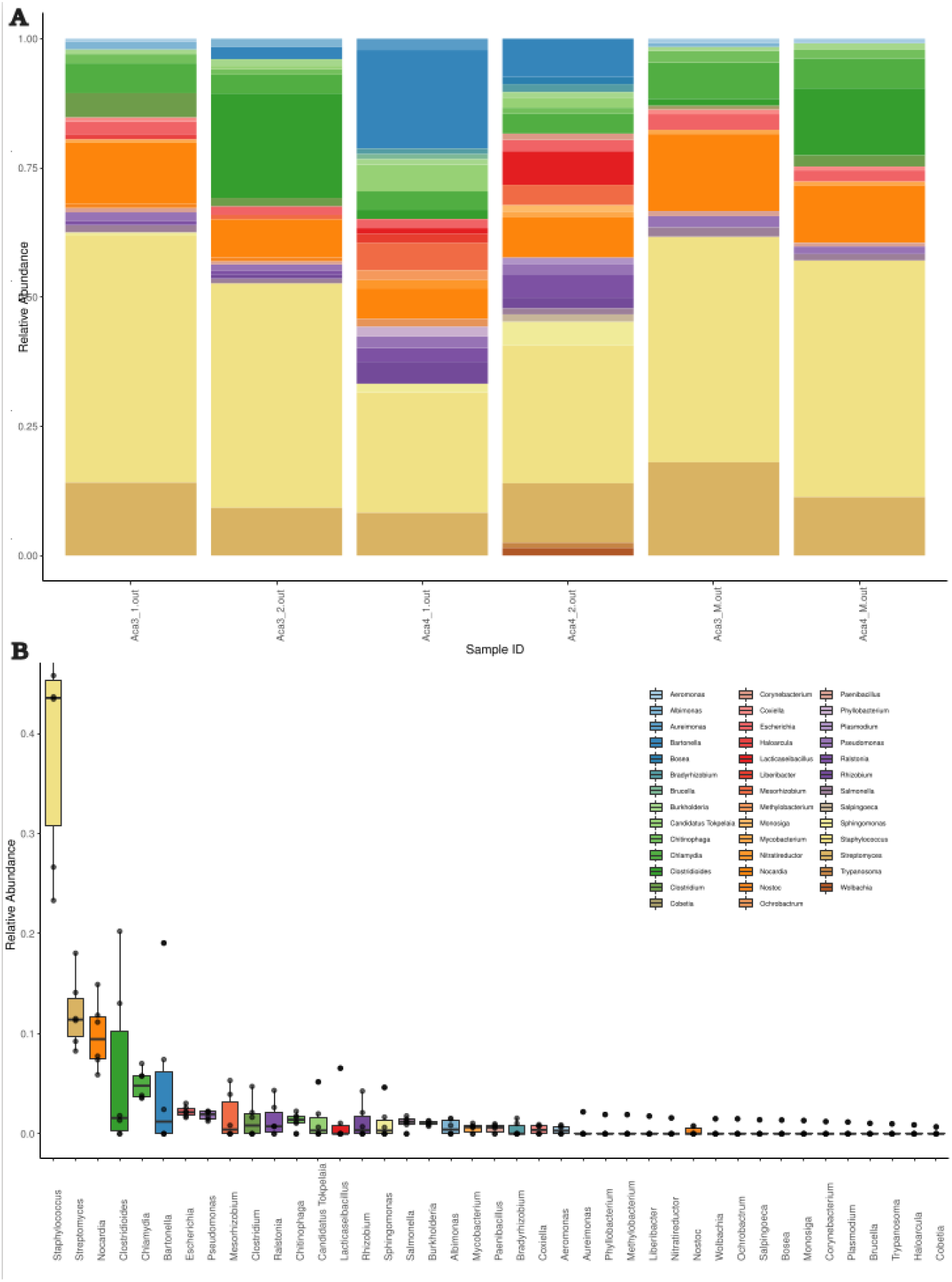
Kaiju results. (A) Stacked bar plot showing bacterial relative abundances (at the genus level) in each mite and saliva sample. The four first bar plots corresponded to the mite samples (namely Aca 3 for one host and Aca 4 for the other host), and the last two samples corresponded to the saliva from both hosts (B) Boxplot summarizing the relative abundance of each genus of bacteria assembled.

**Figure 3.**
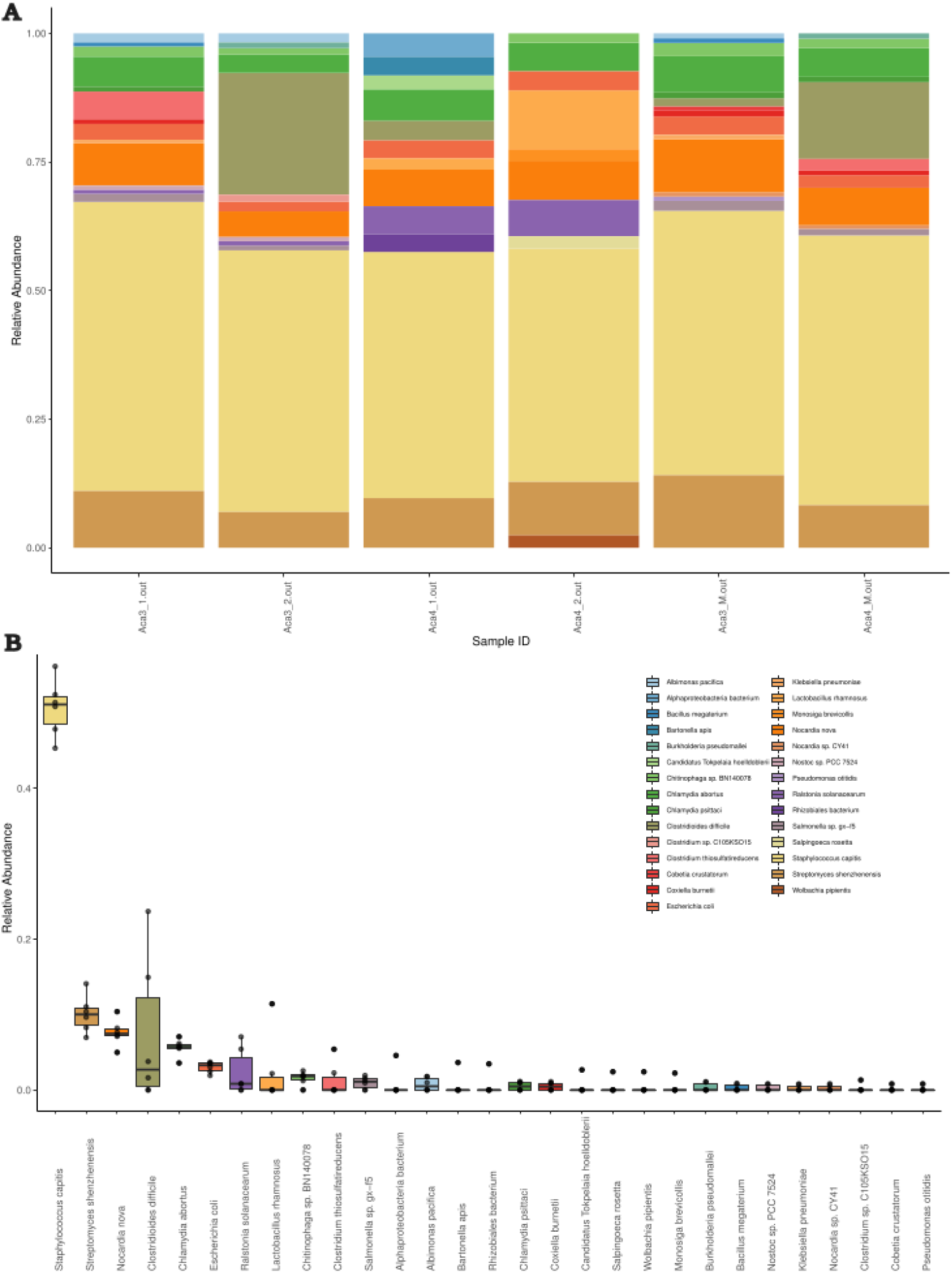
Kaiju results. (A) Stacked bar plot showing bacterial relative abundances (at the species level) in each mite and saliva sample. The four first bar plots corresponded to the mite samples (namely Aca 3 for one host and Aca 4 for the other host), and the last two samples corresponded to the saliva from both hosts (B) Boxplot summarizing the relative abundance of each genus of bacteria assembled.

## 4. Discussion

In this study, we conducted shotgun and genome-resolved metagenomic analyses of a rhinonyssid mite species (*Tinaminyssus melloi*). A preliminary characterization of the microbiomes of this mite species [32] suggested that a potential endosymbiont Alphaproteobacteria (Bartonellaceae family) was associated with this mite species. Here, we expanded the sample size, used contamination controls, and increased sequencing depth to retrieve an almost complete genome of this bacterial taxon. In addition, our comparative analyses of the genome of this bacterium support the endosymbiotic nature of this taxon, which is the first endosymbiont described to date for nasal mites.

The genome of this endosymbiont was evaluated to be 98.1 % complete. Indeed, considering this value is very close to 100, and given that endosymbionts tend to have eroded genomes (i.e., lose some genes when compared to free-living counterparts), it seems likely that the assembly does in fact represent the complete genome, and it has lost some genes from the panel used to evaluate completeness. This MAG was detected in mite samples but not in the saliva samples, thus supporting its exclusive association with the mites. The comparative analyses indicate that this genome was AT-biased, and the genome size was small (see Results) compared to a previous study on endosymbionts of lice [44]. Altogether, these characteristics have been reported as typical of arthropod endosymbionts [46, 47]. Also, endosymbionts are typically thought to complement deficiencies in their host’s diet [48]. We found that this MAG has complete pathways for vitamin B (riboflavin) and vitamin B12 synthesis. In addition, the MAG has complete pathways for essential amino acid synthesis (i.e., lysine), as well as for several non-essential amino acids (i.e., aspartate, glutamate, serine). We also found many fully or partially missing pathways that may be redundant or potentially shared (or possibly complementary) with the mite. Our analyses on the completeness of metabolic pathways show that it has complete pathways for some essential amino acids (e.g., vitamin B, **Table S2** and **Figure S1**) and non-essential amino acids (e.g., aspartate). We also found many fully or partially missing pathways that may be redundant or potentially shared (complementary) with the mite. Overall, these results are congruent with previous studies of hematophagous parasites in which endosymbionts have been reported and that complement deficiencies in host diet (i.e., *Wolbachia*, *Cardinium*; [47]). However, no endosymbiont has been described from an avian nasal mite species to date; thus, further research on the interaction of bacterial endosymbionts and rhinonyssid mites is needed.

The closest relatives to the potential endosymbiotic bacterial species found by MiGA in both databases were species related to *Brucella (Brucella abortus, Brucella microtis, Brucella pinnipedialis,* and *Candidatus Tokpelaia hoelldoblerii*). However, in all cases, the maximum average amino acid identity was lower than 53 %. *Brucella*-like bacteria has infrequently been described for mite species [49, 50]. Taxator-tk [51], which is based on a much more complete database (NCBI nucleotide), assigned this bacterial taxon to the Bartonellaceae family. Mite endosymbionts belonging to the *Bartonella* genus have been found by previous studies [52–54]. Thus, it may be that this endosymbiont belongs to *Bartonella*, but MiGA databases (TypeMat and NCBI Prok) do not have genomes of closely related *Bartonella* taxa. Indeed, MiGA analyses using both databases indicate that this species is likely to be a new species (P < 0.05) but not a new genus (P > 0.05).

The microbiome composition of hematophagous arthropods has received much attention because their bacterial and virus associates could significantly affect the status of disease in vertebrate host species [55]. Here, we found several bacterial species that have been previously reported in other mite species, such as *Staphylococcus*, *Nocardia*, *Clostridium*, *Bartonella*, and *Chlamydia* [56, 57, 58]. One particular example is that of *Staphylococcus* species. We found some taxa from this genus to be present in a high relative abundance, and Staphylococcal species have been widely described as associates of dust and human skin mites [59–63]. Another example is the genus *Streptomyces*, for which some species have been described as endosymbionts in scabies mites [64], in which their role seems to be providing antimicrobial compounds to the mite. We found *Streptomyces* species in high relative abundance. Overall, the role of *Staphylococcus* and *Streptomyces* in bird’s nasal mites is still unknown and should be explored further. On the other hand, despite their high abundance in Kaiju analyses, we did not retrieve any MAG belonging to any of these genera in our genome-resolved metagenomic approach. It may be that the genomes of these bacterial species are more difficult to assemble (e.g., higher content of repetitive regions), and thus, were discarded along the assembly pipeline because they did not meet the completeness/contamination parameters. Further studies using targeted approaches (e.g., MinYS [65]) are needed to evaluate their potential role as an endosymbiont of bird’s nasal mites. Lastly, we also found species from bacterial genera known to contain species that can cause zoonoses, like *Escherichia*, *Nocardia*, and *Salmonella* [66]. The role these bacteria may have in nasal mites is also unknown, and further dedicated studies are required to understand whether they could be harmful for the birds’ host health.

Overall, apart from this study, whole-genome metagenomic data from nasal mites are not available. Further knowledge on this topic is important because nasal mites could cause the transmission of bacteria and viruses to their vertebrate hosts, acting as vectors of disease for birds. Thus, nasal mites could directly affect the health of avian hosts (or others should they switch hosts, [67]) and other species (e.g., humans that may feed upon wild birds and ingest the mites). Furthermore, this group of mites could support the zoonotic biological cycles of some microorganisms in their bird hosts. Therefore, further research on nasal mites’ microbiomes and investigating their role as vectors of diseases in nature is needed.

## Author Contributions

COM, JD, and MR conceived this study. COM and JD analyzed the data. KJ supervised the research. COM wrote the first draft, which was revised by all other authors.

## Data Availability Statement

The high-throughput sequencing data of each sample is available at the Sequence Read Archive SRA under the accession numbers XXX and XXX. The metagenome-assembled genome is available at NCBI (submission ID: XXX).

## Acknowledgments

We thank the “Centro Municipal Zoosanitario of Sevilla” and especially to Francisco Peña Fernández and Rafael Cuadrado Nieto for providing the samples.

