## Supplementary figures and images for "Genome-resolved metagenomic analyses reveal the presence of a bacterial endosymbiont in an avian nasal mite (Rhinonyssidae; Mesostigmata)"

### Figure S1

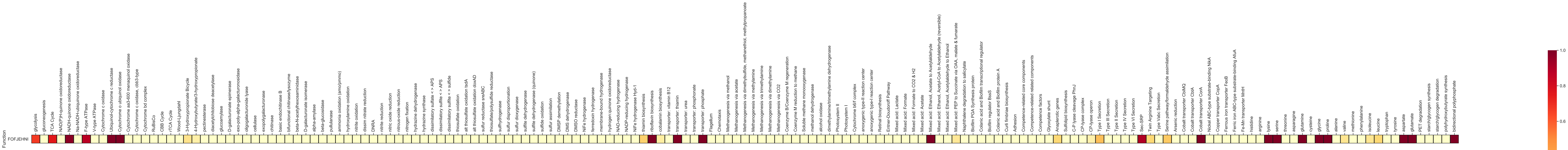


**Figure S1.** Metabolic pathways of the metagenome-assembled genome
