## Supplementary material for "Genome-resolved metagenomic analyses reveal the presence of a bacterial endosymbiont in an avian nasal mite (Rhinonyssidae; Mesostigmata)": Figure S2

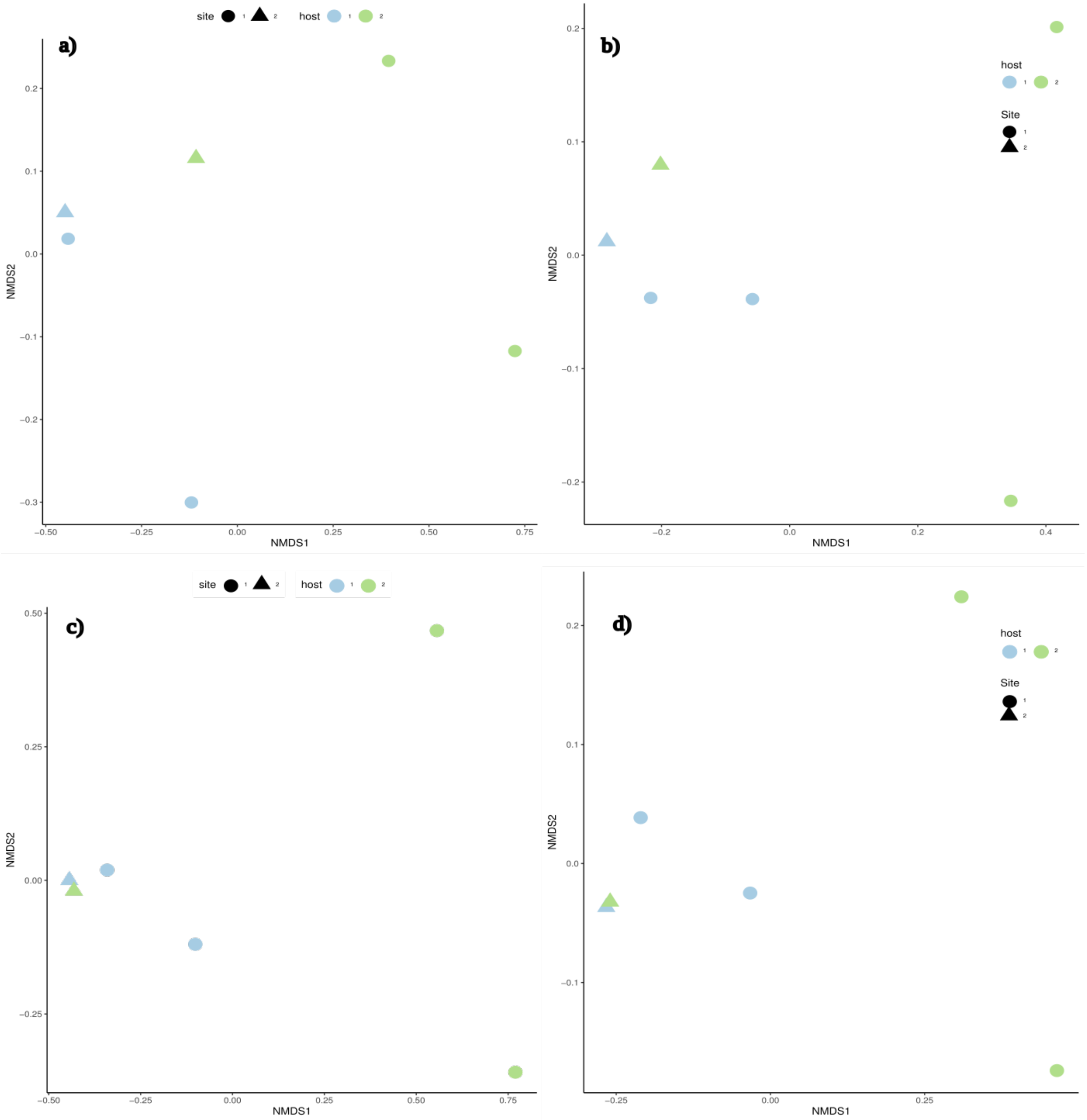


**Figure S2.** NMDS ordinations of mite samples (circle shape) and saliva samples (triangle shape) for each host collapsed to the species level (a and b) and genus level (c and d) based on Bray–Curtis dissimilarity (a and c) matrices and Jaccard distances (b and d).
